# Transplantation of bioengineered lung using decellularized mouse lungs and primary human endothelial cells

**DOI:** 10.1101/2024.08.26.609632

**Authors:** Takaya Suzuki, Tatsuaki Watanabe, Fumiko Tomiyama, Takayasu Ito, Yoshinori Okada

## Abstract

Lung transplantation is a critical treatment for patients with end-stage lung diseases like idiopathic pulmonary fibrosis, but challenges such as donor shortages and post-transplant complications persist. Bioengineered lungs, integrating patient-specific cells into decellularized animal scaffolds, present a promising alternative. Despite progress in using bioengineered lungs in animal models, functionality and structure remain immature. This protocol addresses a critical barrier in organ bioengineering: the need for a cost-effective experimental platform. By using mouse models instead of larger animals like rats or swine, researchers can significantly reduce the resources required for each experiment, accelerating research progress. The protocol outlines a detailed procedure for lung bioengineering using mouse heart-lung blocks and human primary cells, focusing on isolation strategy for the mouse heart-lung block, decellularization, bioreactor setup, perfusion-based organ culture, and orthotopic transplantation of bioengineered lungs. This mouse-scale platform not only reduces experimental costs but also provides a viable framework for optimizing cell types and numbers for recellularization, testing different cell types using histological and molecular methods, and ensuring blood flow post-transplantation. The method holds potential for broad applications, including studying cell interactions in three-dimensional culture conditions, cell-matrix interactions, and ex vivo cancer modeling, thereby advancing the field of organ bioengineering.

**SUMMARY:** This paper describes how to create bioengineered mouse lungs using the decellularization and recellularization methods. It also describes subsequent orthotopic lung transplantation in detail.

## INTRODUCTION

Lung transplantation has been the decisive cure for patients having end-stage lung disease ^1^ such as idiopathic pulmonary fibrosis, where drug treatment is yet ineffective to stop the deterioration of respiratory function. More eligible patients add up to the waiting list every year; however, the number of organ donations from deceased donors has been trailing the increasing number of waiting patients^2,3^. Even after undergoing lung transplantation, quite a few problems would degrade the function of transplanted lungs, including primary organ dysfunction, reactive allogenic syndrome, infections, and so on, which significantly lower the 5-year survival of the lung transplantation recipients^4^.

Several options exist to counter the current problems in organ transplantation, including the utilization of marginal donors^5^, recovery of donor lungs in an ex vivo lung perfusion system^6^, and xenotransplantation using gene-edited swine^7^. These alternatives can expand the pool of donor organs; however, none can entirely address the donor organs’ scarcity, immunogenicity, and functional heterogeneity.

It is far from reality, but bioengineered artificial organs where patient-specific cells are integrated into the decellularized animal organ scaffold are a fascinating potential source of solid organ transplantation f^8^. Several pioneering studies that demonstrated the potential utility of bioengineered lungs have been reported since 2010^9,10^. In these studies, lungs from rats or swine were decellularized by detergents, animal cells or human cells were injected from the trachea or pulmonary vasculature to regenerate the lung tissue in the perfusion-based bioreactor, and some of them were transplanted orthotopically into the animal thoracic cavities^11-15^. However, the function and structure of the bioengineered lungs were premature.

One obstacle to advancing research in organ bioengineering is the lack of a small-scale experimental platform. While rats or swine are the commonly used animals in this field, they require >10^8^ lung cells per lung^16^, which is highly costly to academic labs. If mice are available for organ bioengineering research, we could dramatically reduce the cost of each experiment and speed up the research program. Because the basic architecture of the lung is similar across mammals, the result of mouse-scale experiments can apply to larger animals by simply multiplying the number according to the body size.

The objective of this protocol is to describe the detailed experimental procedure of lung bioengineering using mouse heart-lung blocks and human primary cells^17^.

## PROTOCOL

All experiments followed the Regulations for Animal Experiments and Related Activities at Tohoku University (15th edition), published by Tohoku University (https://www.clag.med.tohoku.ac.jp/clar/en/). This study was approved by the Institutional Animal Care and Use Committee at Tohoku University (#2020AcA-041-01).

1. **DECELLULARIZATION OF MOUSE LUNGS**
  1.1. Preparation of decellularization solutions (1000 mL format in a 1L Pyrex bottle)
    1.1.1. Sterile water: Add 1000 mL of distilled or deionized water to a 1 L format Pyrex bottle. Autoclave (20 min at 121°C).
    1.1.2. 0.1% Triton X100: Add 1 mL of Triton X100 to 1000 mL of sterile water in a 1000 mL format Pyrex bottle. (Optional) Add 10 mL of Penicillin & Streptomycin solution. <u>DO NOT AUTOCLAVE</u>.
    1.1.3. 2% Sodium Deoxycholate: Add 20 g sodium deoxycholate powder to 1000 mL sterile water in a 1000 mL format Pyrex bottle. Close the cap and flip the bottle to solubilize the powder. (Optional) Add 10 mL of Penicillin & Streptomycin solution (Invitrogen). <u>DO NOT AUTOCLAVE</u>.
    1.1.4. 1M NaCl: Add 29.2 g of NaCl to 1000 mL of distilled or deionized water in a 1000 mL format Pyrex bottle. Autoclave.
    1.1.5. DNase I Stock Solution: Dilute at a 10 mg/mL concentration in medium a Medium A. Medium A: Prepare 5 mM CaCl2 (5 mg CaCl2 in 9 mL DI) and dilute ten times with sterile water.
    1.1.6. DNase I Working Solution: Add 33 µL of DNase I Stock Solution to 10 mL of Medium B.
    1.1.7. Medium B: Prepare 10x Medium B (155mg MgSO4 + 220 mg CaCl2 in 100 ml sterile water) and dilute ten times with sterile water.
  1.2. Preparation of the pulmonary artery catheter and the tracheal catheter (Sterile procedure)
    1.2.1. Cut the rat jugular vein catheter at a length of approximately 15 mm.
    1.2.2. Move the collar to the end of the catheter.
    1.2.3. Insert a PinPort into the catheter and attach it to a PinPort injector. Prepared pulmonary catheters are presented in Figure 1A.
    1.2.4. Store the pulmonary artery catheter in 70% ethanol until use.
  1.3. **MOUSE SURGERY FOR HARVESTING HEART-LUNG BLOCK**
    1.3.1. Euthanize a mouse.
    1.3.2. Place the mouse supine position on a surgical table and fix the limbs. Sterilize by spraying 70% ethanol.
    1.3.3. Open the abdominal cavity in the median line to the neck. Split the sternum. Resect the diaphragm to open the thoracic cavities fully. Remove the thymus. (Optional) the inferior vena cava and the right superior vena cava with a 4-0 silk, which prevents the regurgitation of PBS in the following step 1.3.5., thereby enhancing the washout of the blood in the pulmonary vasculature.
    1.3.4. Cut the abdominal aorta for drainage.
    1.3.5. Inject PBS from the right ventricle with a 5 mL sterile syringe with a 27-gauged needle.
    1.3.6. Tape the main pulmonary artery with a 4-0 silk. (Note) The main pulmonary artery and the ascending aorta can be taped together.
    1.3.7. Open a 2 mm window beneath the pulmonary artery valves by cutting the right ventricle wall.
    1.3.8. Insert the pulmonary artery catheter (Step 1.2.) through the window and secure the previously taped 4-0 silk (Figure 1B). (Note) The main pulmonary artery and the ascending aorta can be ligated and secured together.
    1.3.9. Inject 2 mL PBS slowly through the pulmonary artery catheter with a 5 mL sterile syringe. Make sure both lungs slightly expand as PBS is being injected.
    1.3.10. Cannulate trachea and tie in place with 4-0 silk suture (Figure 1C).
    1.3.11. Inject air slowly through the tracheal catheter with an empty 5 mL sterile syringe. Make sure there is no air leakage from the lungs.
    1.3.12. Remove the heart and lung en bloc. (Note) Do not touch the lung surface with any instruments. Any slight touch could result in air leakage.
  1.4. **DECELLULARIZATION OF THE MOUSE LUNGS (3-DAY-Procedure)** All procedures in section 1.4. should be performed in a clean biosafety cabinet. (Day 1)
    1.4.1. Transfer the resected heart-lung block to a plastic Petri dish and incubate the heart-lung blocks in sterile water for 1 hr at 4°C.
    1.4.2. Water rinses – Inject 2 mL sterile water through the trachea catheter three times and 2 mL sterile water through the pulmonary artery catheter with a 5 mL sterile syringe. Pause after each injection to allow the solution to come out as the lung recoils (Figure 1C).
    1.4.3. Inject 2 mL Triton X100 solution into the tracheal catheter and 2 mL into the pulmonary artery catheter.
    1.4.4. Incubate in Triton X100 solution overnight at 4°C. (Day 2)
    1.4.5. Remove the lungs from Triton solution. Water rinses as Step 1.4.2.
    1.4.6. Inject 2 mL of deoxycholate solution into the tracheal catheter and 2 mL into the pulmonary artery catheter.
    1.4.7. Incubate in deoxycholate solution for 24 hrs at 4°C. (Day 3)
    1.4.8. Remove from deoxycholate solution, and perform DI rinses as described in Step 1.4.2.
    1.4.9. Inject 2 mL NaCl solution into the tracheal catheter and 2 mL into the pulmonary artery catheter.
    1.4.10. Incubate in NaCl solution 1 hr at RT.
    1.4.11. Remove from NaCl solution. Water rinses as Step 1.4.2.
    1.4.12. Inject 2 mL DNAse I Working Solution into the tracheal catheter and 2 mL into the pulmonary artery catheter.
    1.4.13. Incubate in DNase Working solution for 1hr at RT.
    1.4.14. Remove from DNase solution. Rinse as in Step 1.4.2, but use PBS solution instead of sterile water. (Note) After decellularization, the heart-lung blocks can be stored in PBS/antibiotics at 4°C for three weeks.
2. CULTURE OF HUMAN PRIMARY CELLS
  2.1. Mix the base media and supplements.
  2.2. Thaw cells in frozen vials in a water bath at 37°C.
  2.3. Mix cells with culture media in a 15 mL conical tube and centrifuge at 500 g.
  2.4. Count cells and subculture cells at an appropriate cell density.
  2.5. Passage cells until reaching the required number of cells.
  2.6. Harvest and resuspend cells at a 0.5-1 × 106 cells/mL density. For complete pulmonary vascular engineering, 3 × 107 HUVECs are optimal.
3. BIOREACTOR SETUP AND PERFUSION ORGAN CULTURE
  3.1. Preparation of an organ chamber and a cell reservoir
    3.1.1. Cut holes in a silicon stopper using a cork borer, as shown in Figure 2A. Each hole number (i-v) in Figure 2A corresponds to the hole numbers in Figure 2B.
    3.1.2. Insert Masterflex/LS Precision Pump Tubing as indicated in Figure 2B.
    3.1.3. Cut holes in a silicon septum of a GL-45 open-top screw cap using a cork borer. Insert an L/S 14 platinum-cured tubing in the hole (Figure 2B and 2C).
    3.1.4. Autoclave the materials above, including the silicon stopper with tubing, the glass canister, the GL-45 screw cap with tubing, a 250 mL Pyrex glass bottle, and an L/S 14 tubing with lure fittings (Tubing B and Tubing C in Figure 3A). Select proper lure fittings to ensure tubing B and C make a loop. (Note) The glass canister is used for an organ chamber, and the 250 ml glass bottle is used for a cell reservoir.
  3.2. Assembly of perfusion-based bioreactor circuit (The following procedures should be performed on a clean bench).
    3.2.1. Add 100 mL of culture media to the glass canister. Then, put the silicon stopper on top of the glass canister.
    3.2.2. Assemble a silicon stopper, a glass canister, a GL-45 screw cap with tubing, a 250 mL Pyrex glass bottle, and an L/S 14 tubing with lure fittings using 3-way stopcocks as described in Figure 3A. Insert a 20-gauged needle in a silicon septum of a GL-45 screw cap.
    3.2.3. Fill the media in Tubings A, B, and C with a 10 mL syringe connected to stopcocks i) and iii). Ensure that there is no air bubble inside the tubing.
    3.2.4. Attach the pulmonary artery catheter of the decellularized heart-lung block via a lure fitting connected to Tubing C. Avoid air bubbles in the catheter or the tubing.
    3.2.5. Add the cell suspension from Step 2.6 to the cell reservoir. (Note) The cell suspension is preferably prepared between Step 3.2.4 and Step 3.2.5. Put a stirring bar in the cell reservoir.
  3.3. Gravity-driven injection of endothelial cells
    3.3.1. Place the cell reservoir containing HUVECs on a magnetic stirrer approximately above the organ chamber (Figure 2D and Figure 3A).
    3.3.2. Turn on the magnetic stirrer.
    3.3.3. Open the stopcock i) and ii) so that the cell suspension can be injected into the decellularized scaffold via Tubing A, Tubing C, and the pulmonary artery catheter. A typical injection rate is 1-2 mL per minute.
    3.3.4. After fully injecting the cell suspension, detach Tubing A from Stopcock ii). Cell count can be performed at this step to measure the cell retention rate in the decellularized scaffold.
  3.4. Perfusion organ culture
    3.4.1. Place the organ chamber in a CO2 incubator.
    3.4.2. Fix Tubing B to EasyLoad III pump head connected to Masterflex pulsatile pump. (Comment: EasyLoad pump heads can be stuck up to 4)
    3.4.3. Close the CO2 incubator’s glass door. Make sure that the tubing is properly placed between the glass door and the rubber seal (Figure 3B).
    3.4.4. Incubate the decellularized scaffold for 3 hours to settle the injected endothelial cells in the scaffold.
    3.4.5. Start the pump at a rate of 6 rpm, which makes 2 ml/min media perfusion using an L/S 14 tubing. Watch the decellularized lung being slightly expanded with the media perfusion.
    3.4.6. Close the incubator door (Figure 3C).
    3.4.7. Media change can be performed through Stopcock iii) by a 50 mL sterile syringe with temporary cessation of pump-driven perfusion. The typical media volume in the organ chamber is 100 mL. The authors usually change half of the media every 2 or 3 days.
    3.4.8. Harvest the recellularized heart-lung block after at least 2-day perfusion organ culture.
4. ORTHOTOPIC TRANSPLANTATION OF THE BIOENGINEERED LUNG
  4.1. Drug preparation
    4.1.1. MMB combination anesthetic agent
    4.1.2. Midazolam (4 mg/kg) (Sandoz, Yamagata, Japan)
    4.1.3. Medetomidine (0.75 mg/kg) (Nippon Zenyaku Kogyo., Fukushima, Japan)
    4.1.4. Butorphanol tartrate (5 mg/kg) (Meiji Seika Pharma, Tokyo, Japan)
    4.1.5. Clinical grade normal saline (0.9% sodium chloride) (Otsuka Pharmaceutical, Naruto, Japan)
    4.1.6. MMB is prepared by mixing these drugs with normal saline.
    4.1.7. Heparin solution (1000 U/ml) (Mochida Pharmaceutical Co. Ltd., Japan)
    4.1.8. Cefazolin Sodium (30mg/kg) (Otsuka Pharmaceutical, Naruto, Japan)
    4.1.9. Cefazolin is prepared by mixing with normal saline.
  4.2. Surgical procedure
    4.2.1. Preparation of cuffs First, lightly rub the three types of angiocatheter with fine sandpaper to make it difficult for the upturned vessels to return. Second, prepare a bronchial cuff from a 20-gauge Tefron angiocatheter 1 mm in length using a #11 scalpel. Before cutting the angiocatheter, use the back of the #11 scalpel to make an indentation in the angiocatheter to tie off the 10-0 nylon. Prepare the pulmonary vein cuff from a 22-gauge Teflon angiocatheter 0.8mm in length and the pulmonary artery (PA) cuff from a 24-gauge Teflon angiocatheter 0.6 mm in length similarly. (comment) This preparation can be done before the day of the experiment.
    4.2.2. Attachment of cuff to bioengineered lung
      4.2.2.1. Place the heart-lung block on a piece of sterile gauze moistened with cold saline, place another piece of dry, clean gauze under it, and put a Petri dish filled with clean ice. Placing the dry gauze prevents the heart-lung block from accidentally freezing.
      4.2.2.2. Place an aneurysm clip to secure the trachea and the gauze (Figure 4A). Place a small piece of gauze on the heart together with the right and left lung separately. Adjust this retraction with the moistened gauze to expose the left hilum as clearly as possible.
      4.2.2.3. Carefully dissect the hilar structures from each other using straight or angled micro forceps, depending on preference (Figure 4B). Dissecting starts from the pulmonary ligament along the vagus nerve. The visceral pleura is incised under the pulmonary vein (PV), and the descending aorta and vagus nerve are moved together through the back of the left pulmonary hilum to the cranial side.
      4.2.2.4. Dissect the left main PA from the pulmonary trunk to the very edge of the left lung (Figure 4C). Then, divide the PA at the level of the pulmonary trunk to obtain adequate length.
      4.2.2.5. Dissect the left PV from the left side of the left atrium to the very edge of the left lung (Figure 4D). Divide the left PV at the level of the left atrium to obtain adequate length.
      4.2.2.6. Suspend the PA cuff just above the PA using the stabilization clamp and insert the PA inside the cuff (Figure 4E). Fold the PA over the cuff), exposing the endothelial surface. Secure around the cuff with a 10-0 nylon tie (Figure 4F). Place the cuff on PV in an identical fashion (Figure 4G and H).
      4.2.2.7. Divide the left bronchus at the level of the carina. Place the cuff on the bronchus in an identical fashion (Figure 4I).
    4.2.3. Procedure for the recipient mouse
      4.2.3.1. Anesthetize the recipient mouse with a mixture of MMB i.p. and intubate by inserting a 20-gauge polyurethane angiocatheter under a microscope. Place the recipient mouse in the right lateral decubitus position and connect the angiochatheter to the respirator. Set the ventilator setting as follows: 2L of O2/min, respiratory rate 120 bpm, 0.5ml tidal volume. Sterilize the chest wall with 70% ethanol. Inject the cephazolin mixture s.c.
      4.2.3.2. Incise the skin with scissors. Cut the subcutaneous tissue and muscles with a cautery. Open the chest through the 3rd intercostal space and place chest retractors. (Note) Although the thoracotomy needs to be large enough for implantation, do not injure the internal mammary artery, which can lead to massive bleeding.
      4.2.3.3. Dissect the pulmonary ligament with a cotton swab and large spring scissors. Place a curved arterial cleanser on the recipient’s left lung so the lung can be retracted easily. With an angled micro-forceps, dissect the mediastinal pleura around the left pulmonary hilum. (Note) The basis of dissecting the pleura is similar to the anatomical lung resection of a human being.
      4.2.3.4. Dissect PA from the bronchus using curved micro forceps from the mediastinum to the very edge of the left lung (Figure 5A). Dissect bronchus from PV in a similar manner. (Note) Dissection can be performed easily when the pleura is dissected at the previous maneuver (4.2.3.3.).
      4.2.3.5. Place a slipknot of 10-0 silk at the base of PA to occlude (Figure 5B). Place a slim angled aneurysmal clip at the base of PV and bronchus (Figure 5C).
      4.2.3.6. Wrap 10-0 nylon around the bronchus, PA, and PV, leaving loose to secure cuffs at the following steps.
      4.2.3.7. Incise the recipient’s PA, bronchus, and PV at the edge of the recipient’s left lung using microspring scissors (Figure 5D). Gently dilate the PA and PV using straight micro forceps. Remove the blood in the PA and PV with saline using a 1ml syringe and a 24 gauge angiocathater. (Note) The Incisions of PA, bronchus, and PV are about one-third of the way around. The straight micro forceps are gentle and suitable for dilation.
      4.2.3.8. Place the bioengineered lung, which is covered with moistened gauze, on top of the recipient’s left lung (Figure 5E). The bioengineered lung must be placed as close to the recipient’s mediastinum as possible.
      4.2.3.9. Inserting the donor PA cuff into the recipient PA (Figure 5F). There will be some stretch on donor PA. Comment. If the PA incision size is appropriate, the donor’s cuff is less likely to be dislodged. Moving the bioengineered lung close to the mediastinum will also prevent the donor’s cuff from being dislodged.
      4.2.3.10. Secure around the cuff with a 10-0 nylon tie (Figure 5G). In a similar manner, insert and secure the donor bronchus (Figure 5H) and PV cuffs (Figure 5I and J).
      4.2.3.11. Remove the slim-angled aneurysmal clip (Figure 5L). It is important to observe that blood is backflowing beyond the PV cuff. Then, remove the silk tie on PA to resume antegrade blood flow to the bioengineered lung.

**Figure 1.**
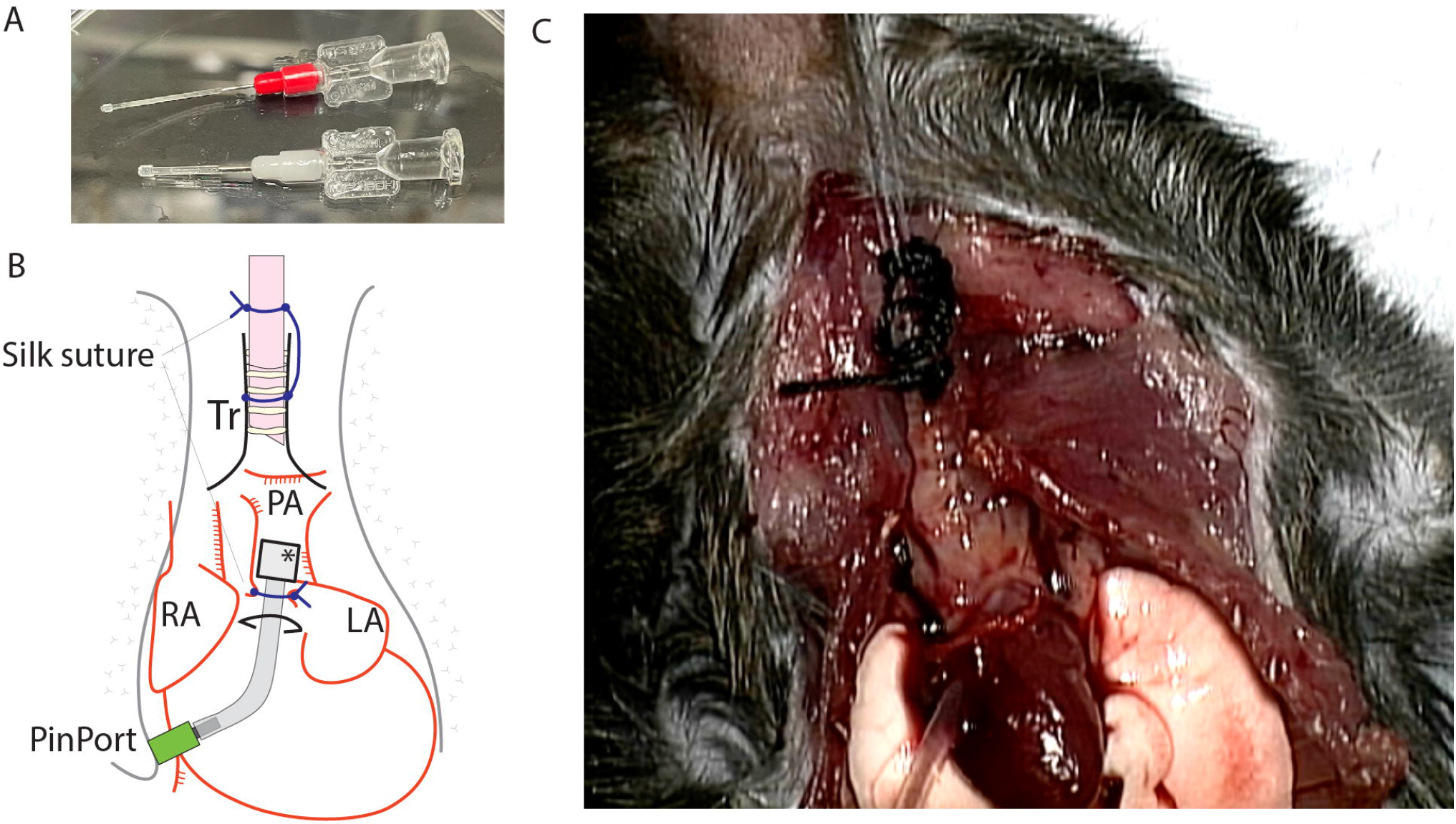
Cannulation of the mouse heart-lung block. A) Prepared pulmonary artery catheters. B) Schema of cannulation. C) Representative image after the completion of cannulation. (Adopted from Tomiyama F., et al., Scientific Reports 2024)

**Figure 2.**
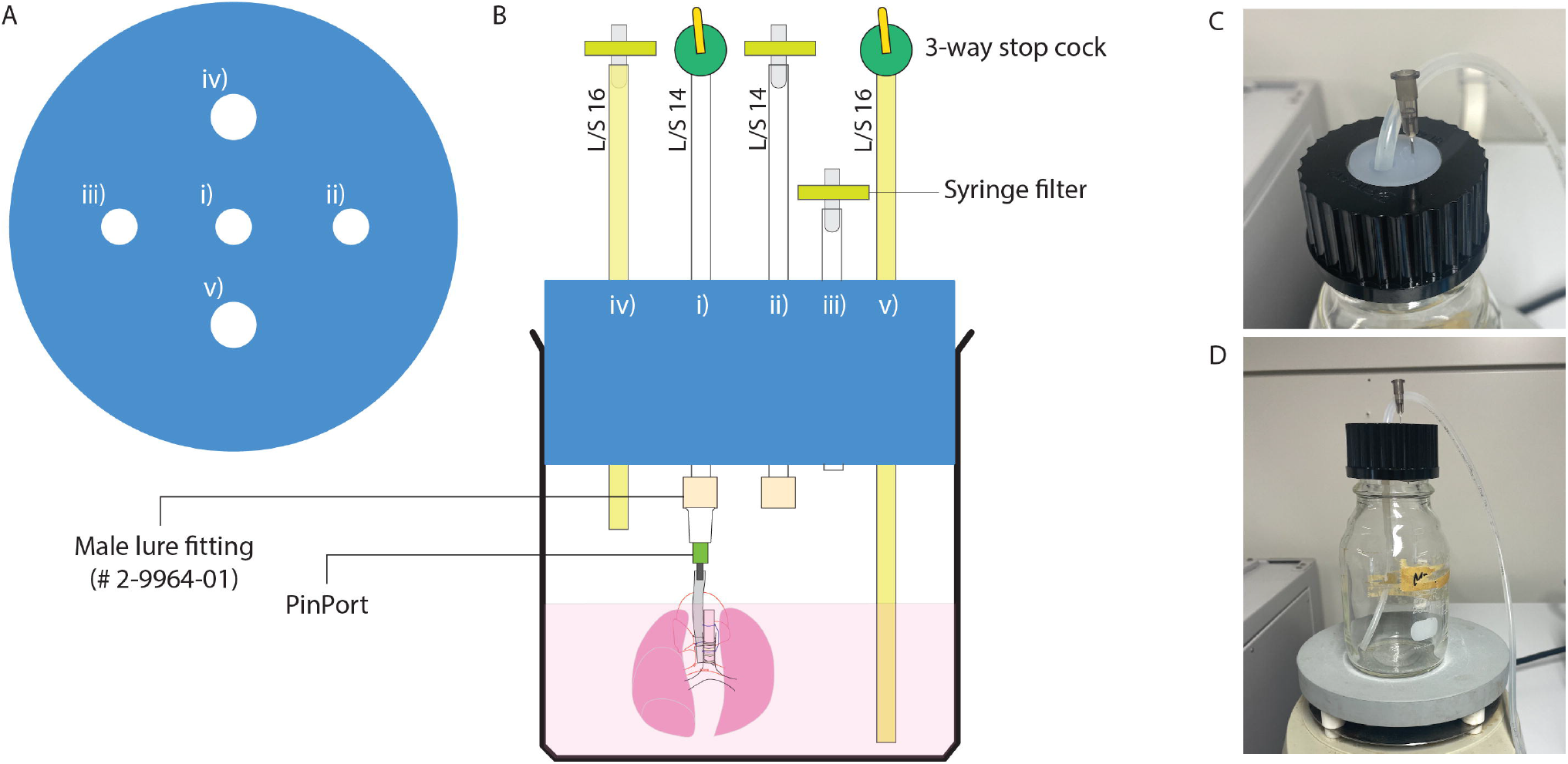
Preparation of organ chamber. A) Holes are cut as described. B) Tubing is inserted as indicated. C) Preparation of the cap for a Pyrex 250 mL glass bottle for the cell reservoir. (Adopted from Tomiyama F., et al., Scientific Reports 2024)

**Figure 3.**
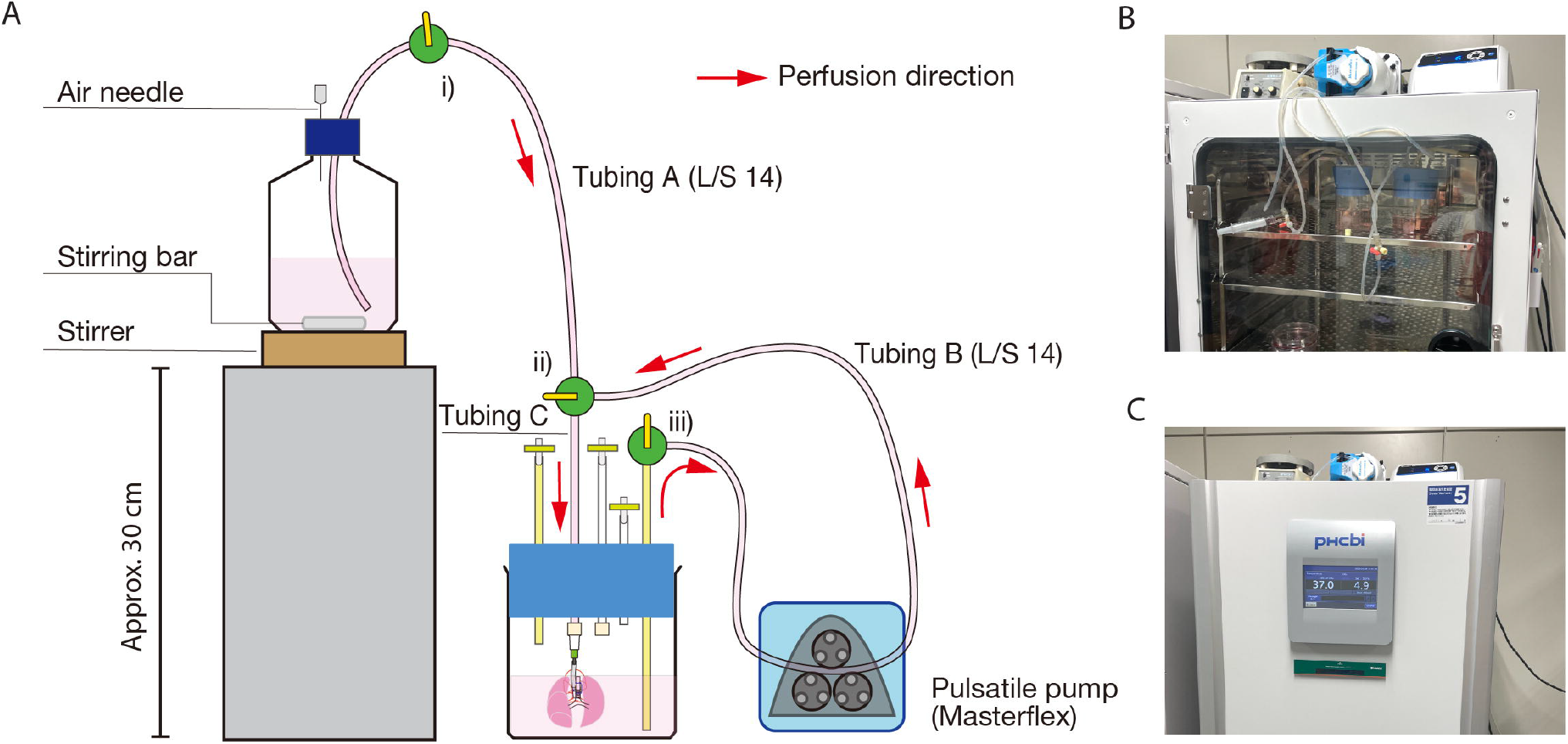
Perfusion-based bioreactor setup. A) Parts and assembly. B) Actual setup. Note that tubing is inserted between a glass door and a rubber seal. C) A Snapshot during the pump-driven perfusion organ culture. (Adopted from Tomiyama F., et al., Scientific Reports 2024)

**Figure 4.**
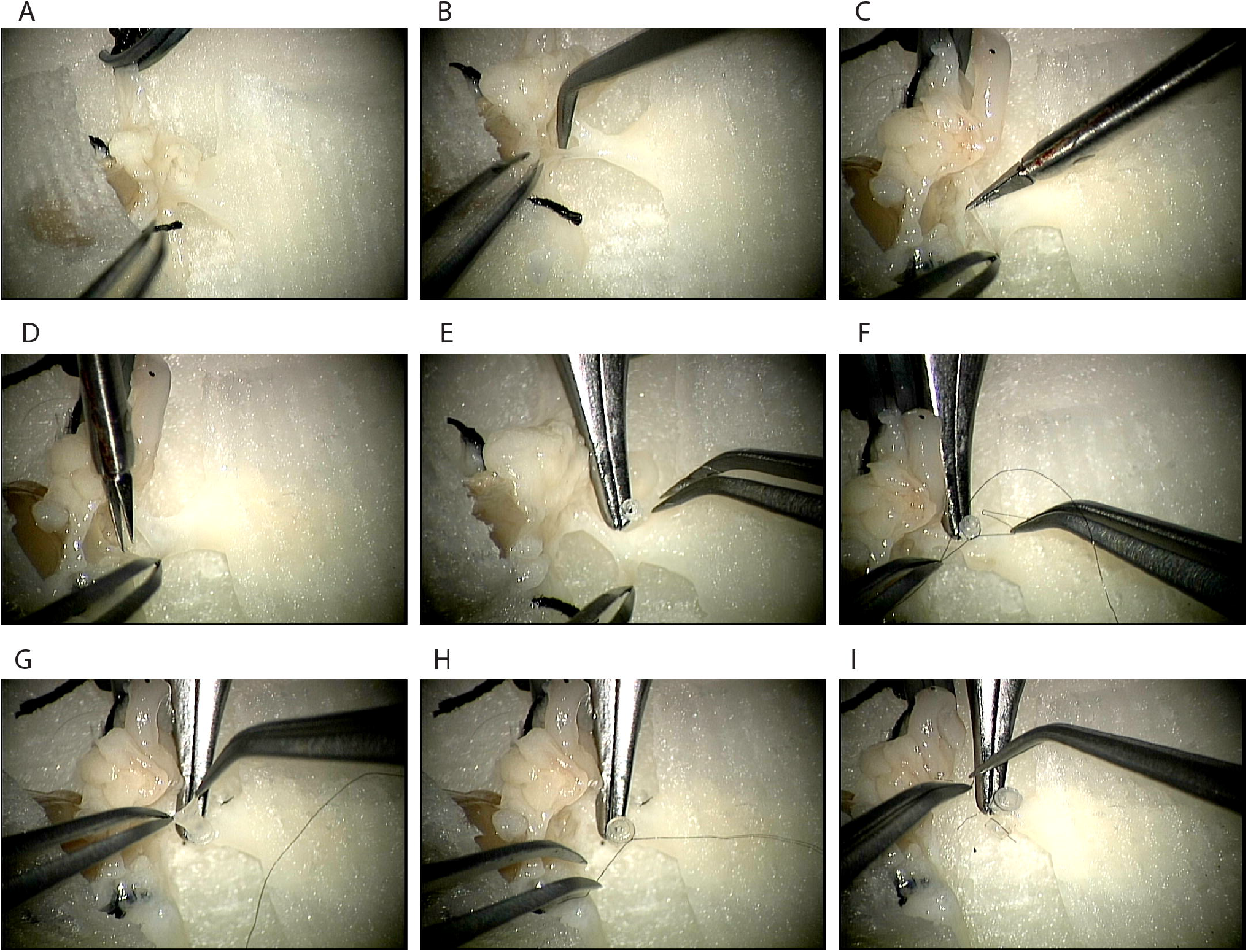
Preparation of the bioengineered lung for transplantation.

**Figure 5.**
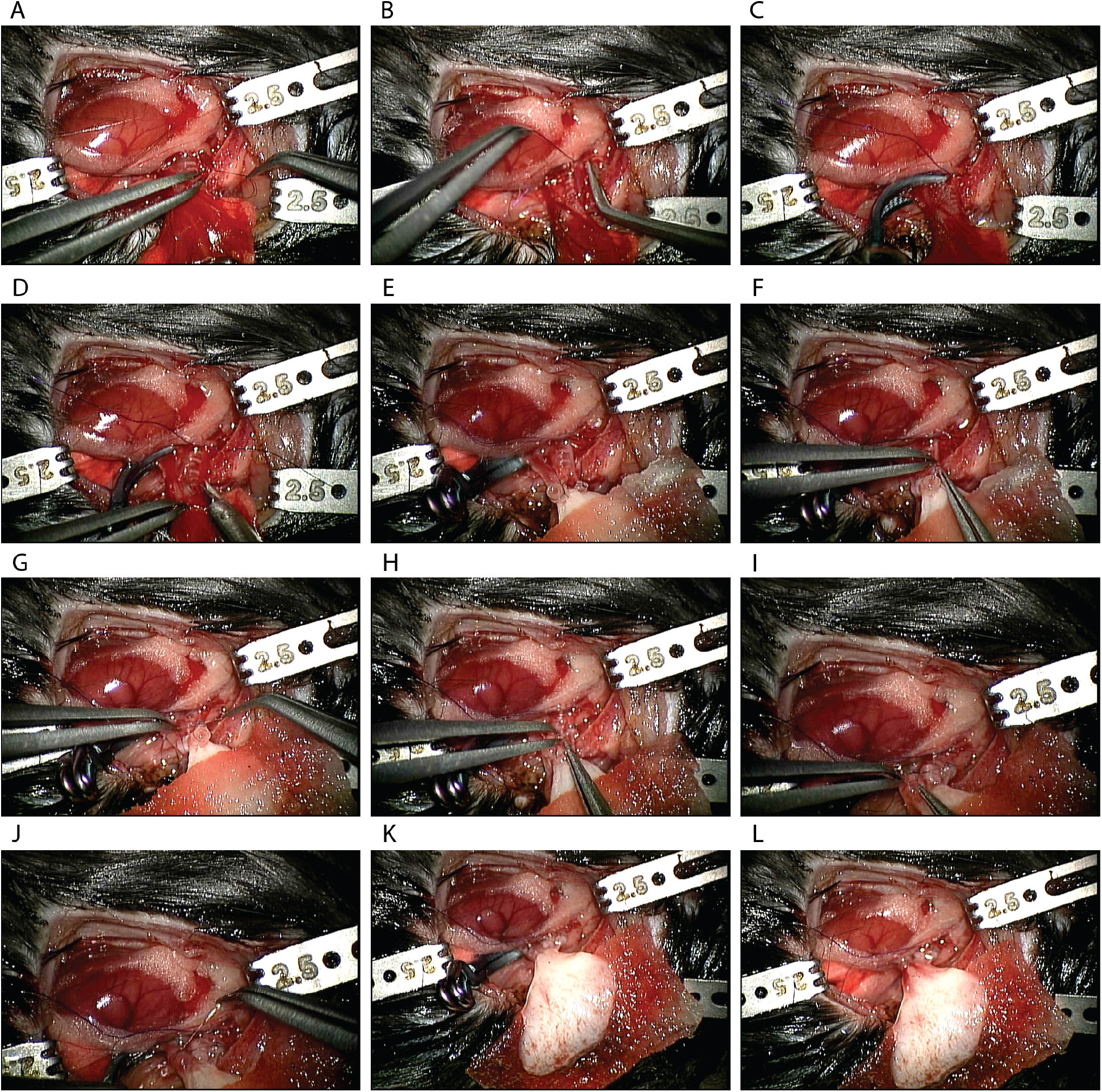
Procedure of orthotopic transplantation of the bioengineered lung.

## REPRESENTATIVE RESULTS

Following the decellularization protocol, mouse lungs are visibly white and translucent (Figure 6A). Cellular components should be entirely removed, but the alveolar structure remains intact in the histological observation (Figure 6B and C).

**Figure 6.**
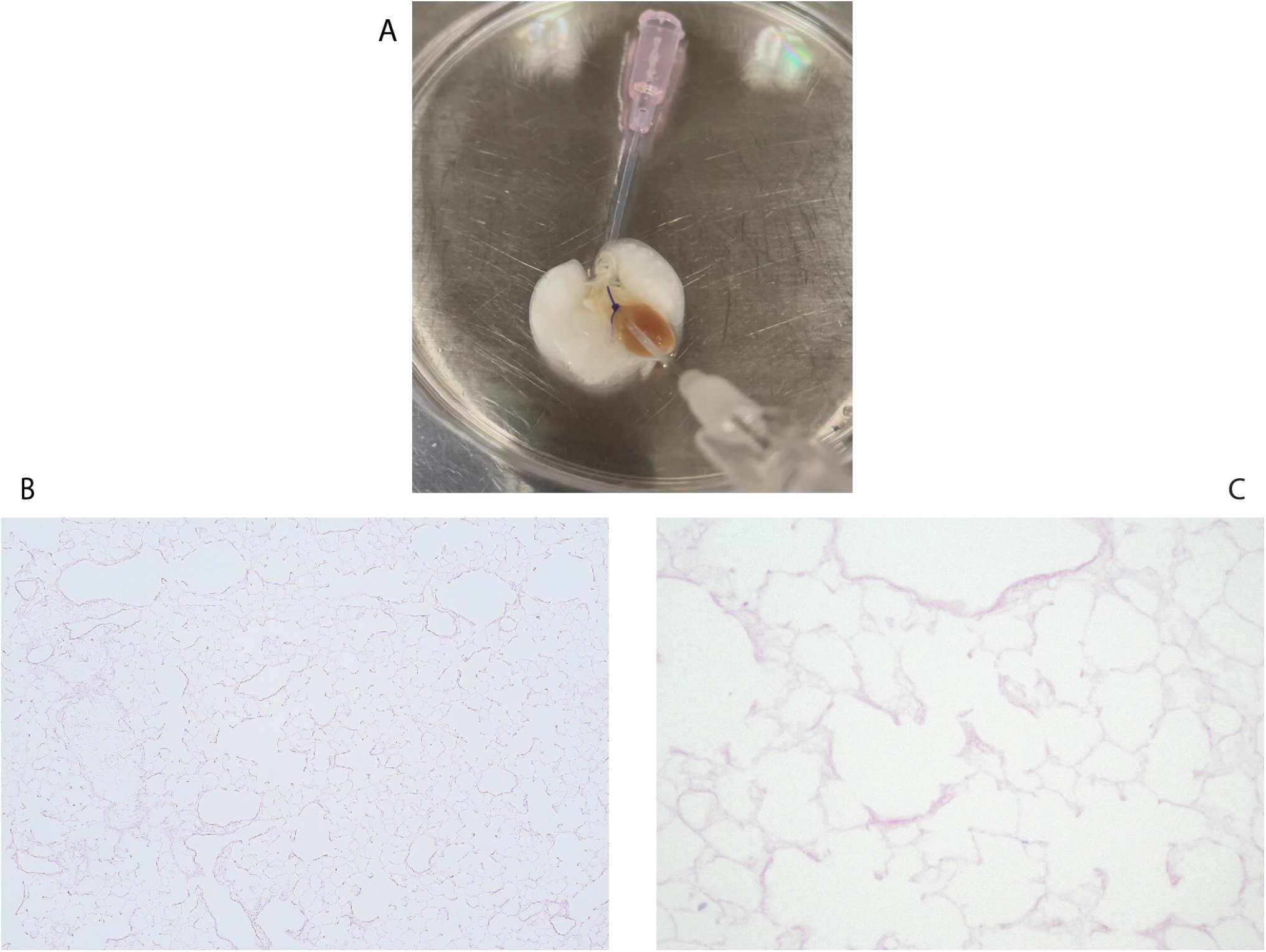
Decellularization of the mouse lung. A) Macroscopic image of the decellularized lung. B) Low-power image of the decellularized lung. C) High-power hematoxylin and eosin (H&E) image of the decellularized lung. Note that there is no visible cellular component.

Recellularized mouse lungs using 3×10^7^ HUVECs and 2-day perfusion-based bioreactor culture show a homogeneous distribution of HUVECs (Figure 7A). HUVECs migrate into the peripheral alveolar area, forming a capillary network (Figure 7B).

**Figure 7.**
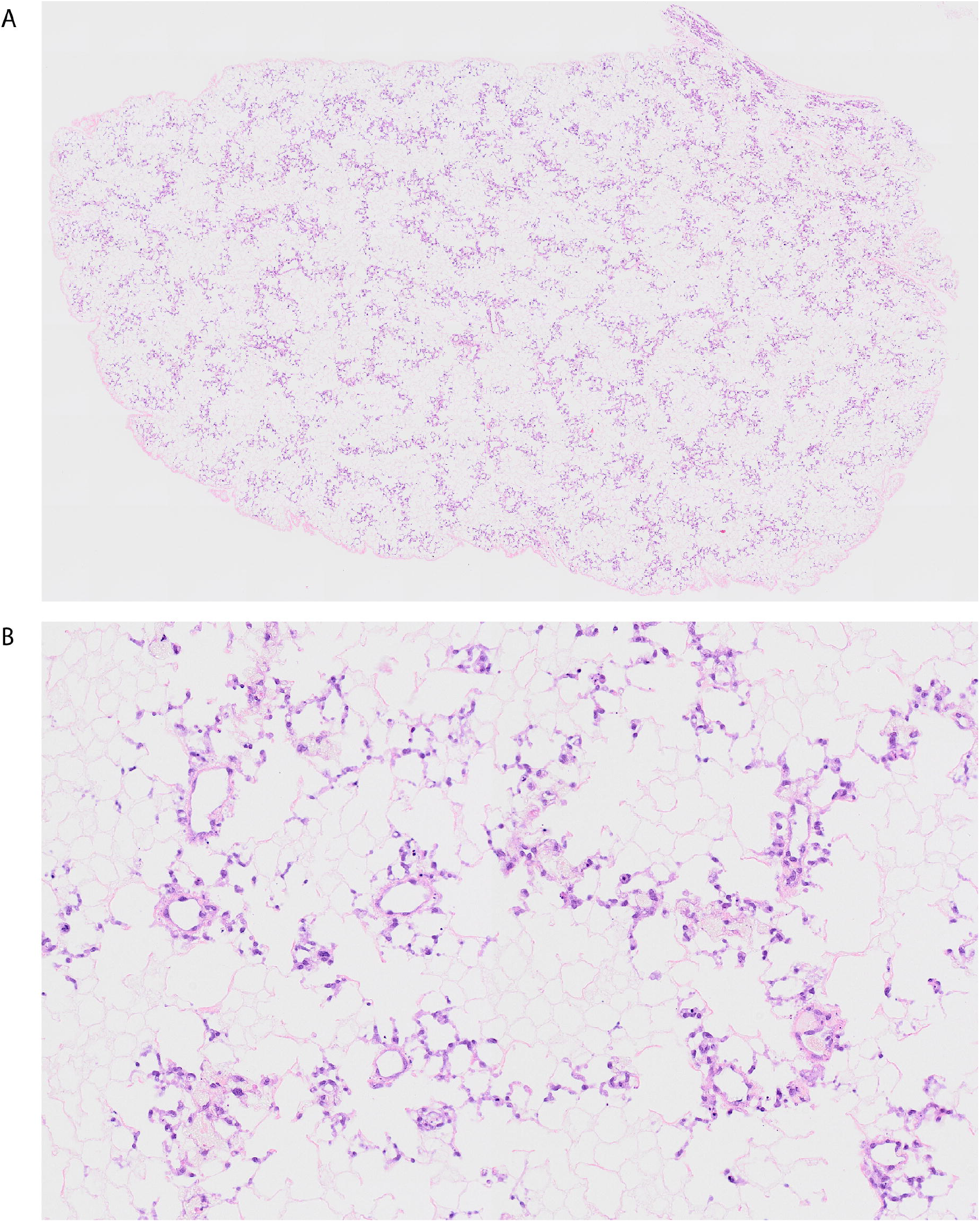
Revascularized mouse lung using HUVECs. A) Low-power H&E image of the revascularized lung. B) High-power H&E image of the revascularized lung

After the orthotopic transplantation and reperfusion of bioengineering lungs, blood flow containing red blood cells is homogeneously observed in the bioengineered lungs (Figure 8A and 8B).

**Figure 8.**
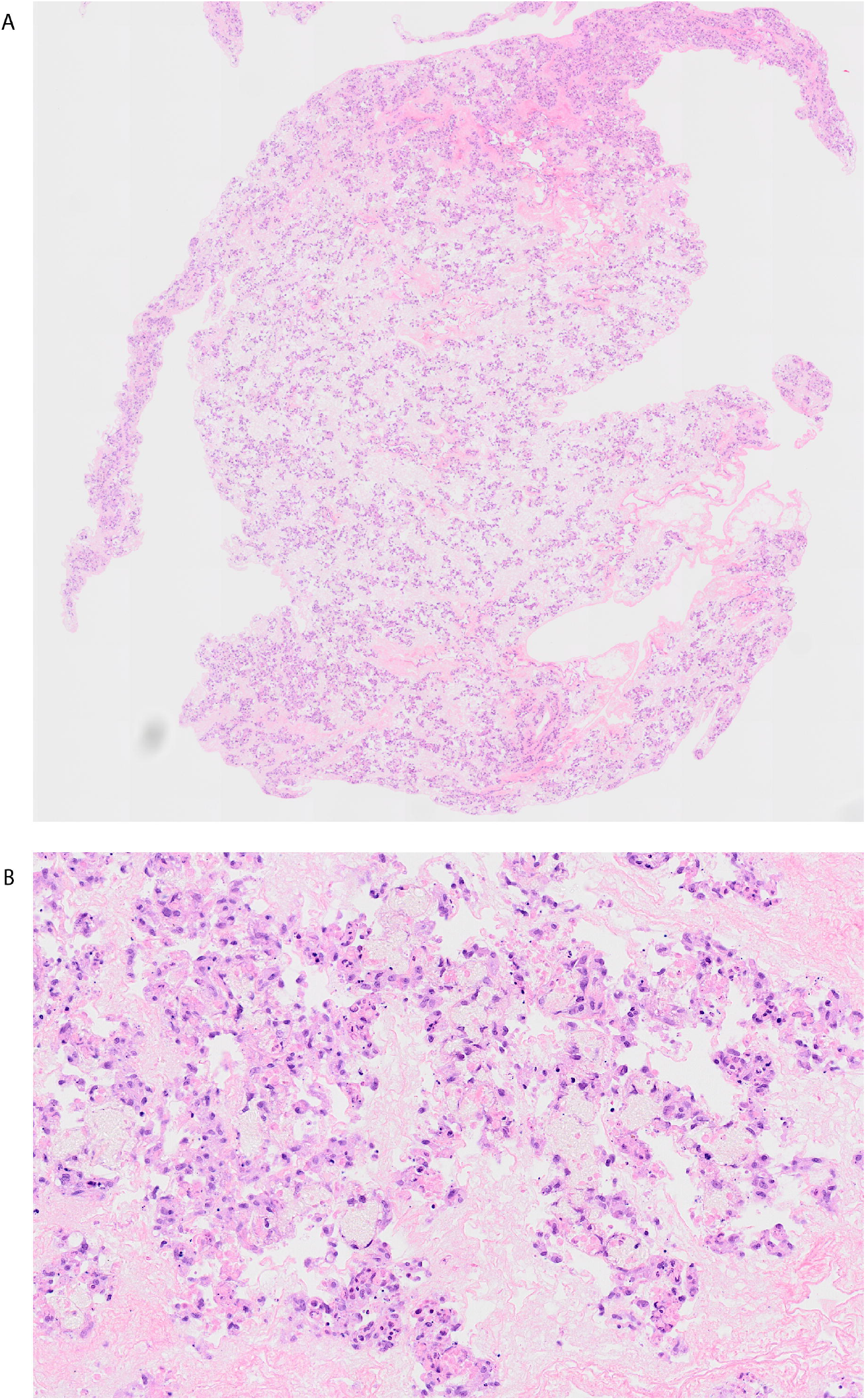
Lung image after transplantation and blood reperfusion. A) Low-power H&E image of the revascularized lung after 10 minutes of reperfusion. B) High-power H&E image of the revascularized lung after 10 minutes of reperfusion.

## DISCUSSION

Organ bioengineering is a demanding enterprise. The costly screening process has been hindering this field’s research and development cycle. By using mice as an experimental platform, space, cells, and media are significantly reduced compared to the previously used rat platform. Although measuring detailed physical parameters such as gas exchange, vascular resistance, or lung compliance has not been achieved yet, the mouse-scale platform we have developed can be used for optimizing cell numbers for recellularization, comparing different types of cells using histological and molecular methods, and confirming blood flow after transplantation.

The critical step of the procedure is inserting and fixation a pulmonary artery catheter. The fixation of the pulmonary artery catheter is only possible by utilizing a small diameter catheter (<3Fr) with a collar at the tip. Because of the fragile nature of the lungs, surgery should be performed entirely with caution. Any metal instruments should not touch the lung surface; otherwise, the lung would suffer significant leakage. Use a cotton swab to maneuver the lungs when necessary.

The decellularization protocol described here is based on previous reports ^18,19^. Other protocols using different detergent sets can be applicable. The heart-lung block should always be treated with caution. Typical incidents during the decellularization procedure include the penetration of the pulmonary artery catheter, come-off of the tracheal catheter, and air leakage. The authors have not experimentally confirmed the integrity of the decellularized scaffold after refrigerated storage in PBS. Still, the authors have not experienced problems using decellularized heart-lung blocks stored in PBS for up to 4 weeks.

Avoiding bacterial contamination is crucial. All glass, PVDF, and silicon parts must be autoclaved before the experiment. The other parts should be used only once. To minimize the risk of bacterial contamination, all procedures should be performed in a clean biosafety cabinet. It is desirable to include antimycotics as well as antibiotics in the media. Frequent media changes during the perfusion increase the risk of contamination. In addition, air bubbles must always be avoided in the tubing. Air bubbles in the tubing are subsequently trapped in the decellularized scaffold, which could blockade media perfusion in the peripheral area and result in heterogeneous cell distribution. Also, endothelial cells should be thoroughly detached by trypsinization or other appropriate cell dissociation media. Cell pellets should be broken well to make a homogeneous single-cell suspension. Too much cell density (e.g.,> 2 million cells/mL) could promote the formation of cell clumps, which could result in embolism in the proximal vasculature.

The basis of the bioengineered lung graft preparation is similar to a regular mouse lung transplantation^20,21^. The engineered lung tissue is not as fragile as a regular lung graft. The challenge is that the lung tissue, including the hilum structure, is entirely white or almost transparent. A precise understanding of local anatomy is indispensable for successful transplantation. The stable technique should be earned using native lungs.

Although it was not tested in the current method, whole lung bioengineering using endothelial and epithelial cells should be easy, considering the difficulties in pulmonary vascular engineering described here. Also, this mouse-scale platform can be expanded to other fields of research, such as the investigation of cellular interaction in 3D culture conditions, cell-matrix interaction, ex-vivo cancer modeling, and so on. In summary, this method provides a reasonable and robust lung bioengineering platform.

## ACKNOWLEDGMENTS

This study was financially supported by the Grant-in-Aid for Scientific Research / KAKENHI (C) #20K09174, #23K08308, the Fund for the Promotion of Joint International Research (Fostering Joint International Research (B)) #22KK0132 for TS, JSPS KAKENHI Grant Number 21K08877 for TW, Leave a Nest Grant Ikeda-Rika award for FT, and the Grant-in-Aid for JSPS Fellows #21J21515 for FT. We greatly appreciate Ms. Maiko Ueda, technical staff in the Biomedical Research Core of Tohoku University Graduate School of Medicine, for her intensive work in histological observation. We also appreciate the technical advice of Ms. Yumi Yoshida and Mr. Koji Kaji in the Center of Research Instruments at IDAC, Tohoku University, for their image processing support.

